# *CACTI: An in-silico* drug-target prediction tool through the integration of chemogenomic data and clustering analysis

**DOI:** 10.1101/2024.02.23.581805

**Authors:** Karla P. Godinez-Macias, Elizabeth A. Winzeler

## Abstract

It is well-accepted that knowledge of a small molecule’s target can accelerate optimization. Although chemogenomic databases are helpful resources for predicting or finding compound interaction partners, they tend to be limited and poorly annotated. Furthermore, unlike genes, compound identifiers are often not standardized, and many synonyms may exist, especially in the biological literature, making batch analysis of compounds difficult. Here, we constructed an open-source annotation and target prediction tool that explores some of the largest chemical and biological databases, mining these for both common name, synonyms, and structurally similar molecules. We used this Chemical Analysis and Clustering for Target Identification (CACTI) tool to analyze the Pathogen Box collection, an open-source set of 400 drug-like compounds active against a variety of microbial pathogens. Our analysis resulted in 4,315 new synonyms, 35,963 pieces of new information and target prediction hints for 58 members.

**Scientific Contribution:** With the employment of this tool, a comprehensive report with known evidence, close analogs and drug-target prediction can be obtained for large-scale chemical libraries that will facilitate their evaluation and future target validation and optimization efforts.

## Introduction

Understanding how drugs interact with its biological target and whether it results in the desired phenotypic response is a critical step in drug discovery(1–3). One area where this is particularly true is for infectious diseases where many of the hits that are advanced for drug discovery come from phenotypic, organismal screens as described in a recent review(4). For malaria parasites, tens of thousands of compounds with parasite killing activity have been placed in the public domain. This has led to medicinal chemistry programs and molecules entering clinical trials(4, 5). While understanding the mechanism of action or knowing the target is not strictly essential for a compound series to advance, knowing the target and ideally having a crystal structure for the target can make subsequent medicinal chemistry optimization much more efficient: fewer compounds need to be made and evaluated, and biochemical assays are typically more robust and lower cost. Furthermore, screening and assay costs can be many orders of magnitude lower for a biochemical target relative to whole organism work.

Because of the huge-cost savings that can come from knowing a target, substantial effort is often invested into target discovery. Biochemical and genetic approaches (e.g. in vitro evolution) are two examples of methods used for target identification. However, these tend to be lengthy and resource consuming(4, 5). For malaria parasites, *in vitro* evolution has been a successful method but can often take six months or more(4). It thus makes sense to accomplish as much computationally as possible. For example, molecular docking is commonly pursued to predict small molecule ligand binding to key therapeutic proteins prior to biochemical or phenotypic assays, and though it tends to have a high false positive rate(6), drug targets for some antimalarial inhibitors have been identified through this approach(7–9). Comparative *in silico* approaches also exist for target identification, for example the tool TargetHunter by Wang *et al.*(10) is a web prediction approach that incorporates analog bioactivity data from ChEMBL(11), similarly to Chemmine, an online resource by Backman *et al.*(12) that predicts based on similar records in PubChem database(13). A constraint for these resources is the limit to one molecule per search and one chemogenomic database. Modern high-throughput screens may involve millions of compounds and thousands of primary hits. Having high-throughput methods to rapidly assess many hits can allow prioritization of compounds for resupply and resynthesis, which is particularly important for academic researchers that may have limited access to compound management groups or limited medicinal chemistry capacity. For example, in malaria parasites, many dihydrofolate reductase inhibitors have been compromised by widespread resistance(14). It thus makes sense to identify phenotypic hits acting against DHFR prior to compound resupply.

To address the need of an automated multicompound target prediction tool, we constructed a pipeline named CACTI (Chemical Analysis and Target Identification), to predict biological targets with single or bulk queries, integrating data not only from major databases such as ChEMBL and PubChem, but also from ligand-based databases (e.g. BindingDB(15)), as well as scientific and patent evidence (e.g. PubMed(16) and SureChEMBL(17)). Furthermore, to account for the difference in molecule identifier across databases, we implemented a cross-reference method to map a given identifier based on chemical similarity scores and other known identifiers, or synonyms, in the expanded search. With this process, we are able to provide a comprehensive large-scale study report incorporating all available identifiers for each small molecule, its close analogs, and available bioactivity data and/or mechanism of action (MoA) if known.

## Materials and methods

### Chemogenomic database selection and accessing

A common first step to evaluate a small molecule as a potential therapeutic, includes the exploration of chemical and biological space to identify known information and potential target. In contrast to most available tools that focus on one database for target-predictions(10, 18–20) we explored multiple chemogenomic databases as knowledge source. The solely criterion for database selection was the availability of REST API, a data request protocol for query and transfer. For chemical- and experimental-based information sources, we selected the well-established ChEMBL and PubChem databases. BindingDB was selected for protein-ligand evidence and, EMBL-EBI and PubMed as sources for comprehensive searches (**Table 1**). Although each bears unique data, we observed some overlap in data content, especially for literature evidence. However, it is important to note the indexing type and database specification prior to integration, as not every record is curated, and query fields could have different meaning. For example, experimental data in PubChem could be from biological assays (dose-response, phenotypic, or target-based) or biochemical assay (profiling or binding), while BindingDB refers to binding affinities assays solely.

**Table 1.** Approximate number of records by entity type for selected chemogenomic and literature databases.

| Database | Chemical Data | Experimental Data | Literature | Patent | Compound-Pathway/Gene |
| --- | --- | --- | --- | --- | --- |
| ChEMBL | 22,399,743 | 20,334,684 | 88,630 |  | 15,398 |
| SureChEMBL |  |  |  | > 17 M |  |
| PubChem | 115,036,406 | 304,160,942 | 38,884,316 | 43,059,414 | Pathway: 240,631 |
| Binding DB | 1,164,246 | 2,707,699 | 39,818 |  | 9,023 |
| EMBL-EBI |  |  | 34,135,964 |  |  |
| PubMed |  |  | ~ 34 M |  |  |

After database selection, we created a custom function to access and retrieve data from each source. Following the corresponding REST API web services, we constructed an html query link with the database domain + web service architecture + data of interest + parameters. Specifically, we used chembl_id_lookup, mechanism, document, compound_record and similarity data functions from ChEMBL. For PubChem we used compound’s data SMILES, name, CID, PubMedID, PatentID, fastsimilarity_2d, synonyms and MolecularWeight functions. PubMed records were found using the PMID and PMC xref from PubChem, and europepmc/search function from EMBL-EBI server. The command, GetTargetByCompound, was used to access BindingDB database. For exact URL query construction please refer to the database web-services(15–17, 21, 22).

### Querying and standardization of chemogenomic databases using SMILES

We reasoned that, to investigate a set of query compounds, the first step was to explore hidden patterns among the various chemogenomic databases containing large datasets of small molecules and their bioactivity. However, these patterns are difficult to reveal when searching across databases, mainly due to the lack of indexing standardization and the existence of multiple equivalent representations of a chemical structure (SMILES(23)). For example, ethanol can be encoded with SMILES OCC, CCO and C(O)C. Furthermore, in ChEMBL the SMILES CCO corresponds to CHEMBL545 but C(O)C is not a valid query, and vice versa the notation C(O)C corresponds to CID702 in PubChem while CCO is not valid. Nevertheless, to test the hypothesis that using SMILES as identifier is sufficient for mapping the chemical space, we used the provided query SMILES as first input (**Fig. S1A**). To reveal the index identity of each query, we used the custom functions to cross-reference across databases using 100% similarity match to ensure the identity of each query.

Due to submission discrepancies across repositories, query identifiers were combined in a “synonym” (common names) column and filtered if they were 1) numerical with no indication of external source origin, 2) IUPAC name from an unreliable source, or 3) duplicated when converted to upper case and removal of special characters such as “:”, or “–”. The remaining synonyms were used to retrieve scientific evidence, patent evidence and additional useful information associated with the compound of interest. We reasoned that expanding the search to include these identifiers would provide more exhaustive research on the query compounds, improving the target-prediction steps when using the additional pieces of evidence. With this filtering, invalid or duplicated query records were removed and bioactivity data, naming synonyms, scholarly evidence, and chemical information across selected chemogenomic databases was integrated.

### Chemical comparison through similarity calculations

In addition to mining with SMILES and common names, we expanded the search to include closely related analogs. (**Fig. S1B**). We first used RDKIT v.2018(24) to convert the query SMILES to a canonical form, generating a unique notation to be queried and compared with equal features. The standardized format (canonical) was used to identify analogs by transforming them to a binary representation called fingerprints, which allows chemical similarity calculations. These similarities were computed using the Tanimoto coefficient(25), *T*, which measures the degree of similarity between a query and target structure as follows:

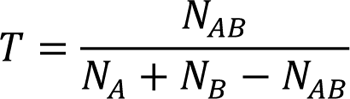

Where *A* represents the query fingerprint and *B* the target fingerprint, *N_A_* and *N_B_* represent the number of “1-bit” in *A* and *B* respectively and *N_AB_* represents the “1-bit” shared in both. To ensure consistency, analogs were transformed to the same canonical form as the query and filtered if the *T* score was below 80% threshold (score ranging from 0-1 transformed to percentage). This conversion allowed us to further reduce discrepancies between SMILES conversion while identifying chemical analogs.

Although a target may have been reported for a given compound, the query compound might be similar but not identical and our goal was to capture these associations. To identify scaffold families, we constructed a similarity network (including the query dataset and searched analogs) by assigning nodes to chemical entities and edges between entities with a Tanimoto coefficient above 80%. With this network we were able to identify query drug-targets from analogs with known mechanisms of action or gene target, elucidating relationships that could be obscured otherwise.

### Drug target identification

In order to predict drug-targets we acquired and incorporated several large datasets with annotated compound-target pairs and created a module to search these sets for compounds closely related to query molecules (**Fig. S1C**). We used the Novartis Chemogenetic Library(26) assembly consisting of 4,185 compounds annotated with their primary mammalian gene target, an in-house consolidated file of 163 validated antimalarials with their drug target originated from phenotypic screens, a collection of 218 licensed well-validated therapeutic drug extracted from well-established databases such as IUPHAR/BPS pharmacology database(27) or FDA approved drugs list (28), and a set of 157 known antibacterial inhibitors(29) from which seven (azithromycin, mupirocin, dapsone, sulfalene, sulfadiazine, triclosan, sulfamethoxazole) overlap with the antimalarial set. The assembled list can be accessed at https://github.com/winzeler-lab/CACTI metadata folder.

To reduce biases when comparing large datasets with molecules of wide mass range, we implemented a method to partition the query and target dataset by molecular weight (default cutoff of 500 Da) and convert each into fingerprints for subsequent similarity calculations. We then implemented a function to allow selection of fingerprint conversion algorithm (RDKFingerprint or GetMorganFingerprintAsBitVect) and selection of binary vector size, to prevent biased structure pattern identifications. Once fingerprints were calculated, we created a NxM matrix, where N and M is the number of fingerprints in the dataset and calculated Tanimoto coefficients for every pair, followed by clustering molecules based on their score (default 0.8).

### CACTI, a large-scale drug-target prediction tool

As a final step we combined the chemogenomic database querying approach and the cheminformatic similarity and clustering approaches to construct a prediction pipeline (**Fig. 1**). CACTI is accessible at https://github.com/winzeler-lab/CACTI. Shortly, we built the customizable pipeline with python version 2.8 currently executable with UNIX, allowing for single or bulk queries and the selection of analyses and parameters to be performed. After submission validation of the query SMILES set (**Fig. 1A**), the first step in the pipeline was to use the chemical and biological space exploration method to obtain the multiple identifiers from which a single compound is known, including IUPAC and common name(s), as well as peer-reviewed publications, deposited datasets, and patents associated to each common name (**Fig. 1B**). To identify analogs in the public domain that might have a known target, we then retrieved the list of similar scaffolds to the ones of interest (**Fig. 1C**) with at least 80% similarity. This step allowed us to increase the likelihood of accurately predicting a drug-target pair, or providing a prediction starting point.

**Figure 1.**
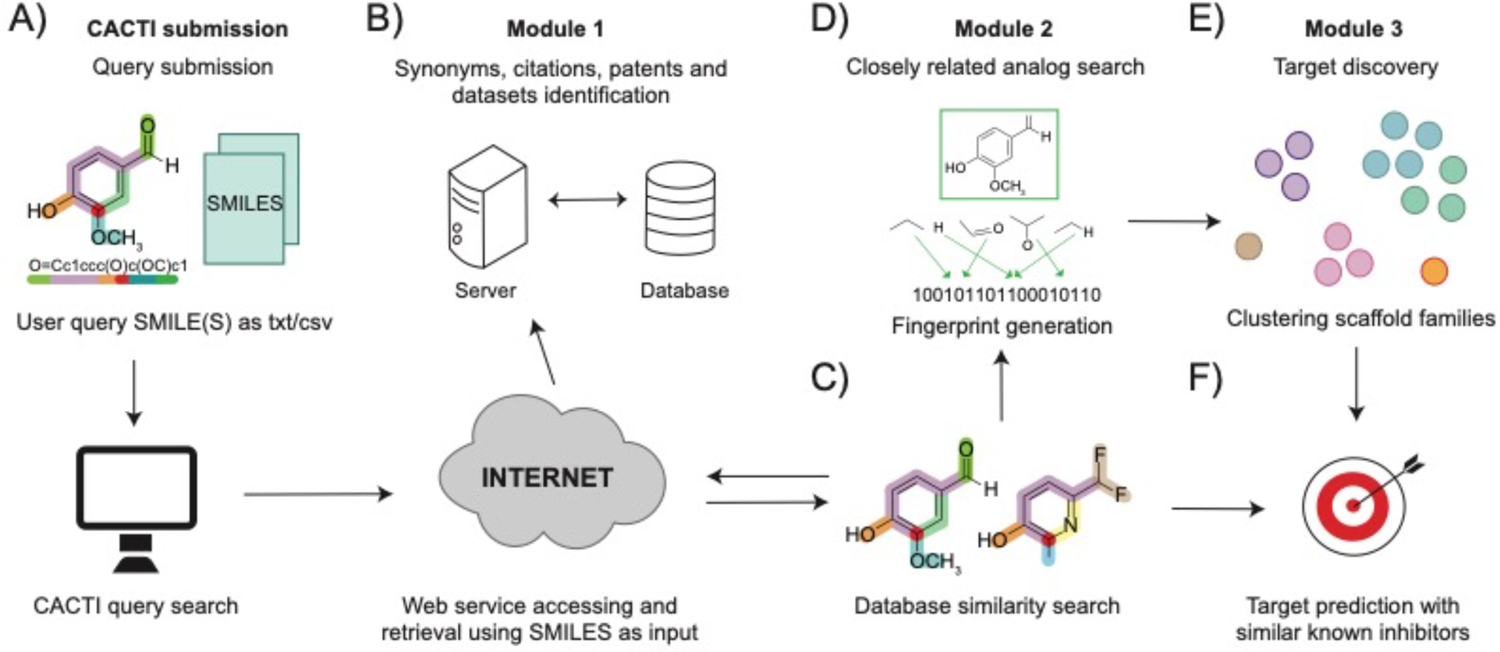
CACTI drug target prediction workflow. Pipeline workflow steps are illustrated by modules where **A)** represents the user SMILES set as input in a tabular or comma delimited, **B)** refers to the database querying and retrieving steps using SMILES to identify synonyms, citation evidence, patents, membership of datasets and additional information for target prediction. Module 2 refers to C) the identification of close analogs in chemogenomic databases to the query set, and **D)** transformation from SMILES to binary fingerprint for scaffold comparison. In module 3, **E)** clustering chemical entities based on similarity measures is performed and **F)** target predictions are completed combining information from previous modules and validated known-target datasets.

To better understand the query molecules and relationships within the dataset and public domain using the acquired genomic and bioactivity annotations, the next step in the pipeline includes the chemical comparison methods. SMILES for the query set and analogs are transformed to binary fingerprints (**Fig. 1D**) to allow comparisons between them, and clustered together to find core scaffolds that are similar to at least 80% to each other (**Fig. 1E**). Although this step seems redundant after querying for similar structures in **Figure 1C**, performing this extra validation was valuable to discover if core scaffolds from the analog subset is shared by two or more query compounds. Lastly, to predict a drug-target under the molecular similarity principal (**Fig. 1F**), scholar references, known mechanisms of action or gene targets, and bioactivity data associated to compound members within each cluster are compared.

## Results

To facilitate the exploration of the chemical and biological space for large datasets, and to provide a better understanding of chemical structures and their impact on a biological system, we constructed a drug-target prediction tool using data mining and chemoinformatic techniques. Our overall objective was to automate the tedious molecule-by-molecule searching that occurs when hits from a high throughput phenotypic screen are initially evaluated and to perform this search in a comprehensive, unbiased fashion, looking for information not only on the query molecules but on closely related analogs. Because we noted that molecules might have different common names in the literature, we also sought to systematically uncover synonyms. For example, the new imidazolopiperazine antimalarial named ganaplacide that is in phase III clinical trials has been known as KAF156, and GNF156 and substantial work has been published on the closely related scaffold named GNF179 which differs from KAF156 by a single halogen substitution. Our motivation was thus to provide a tool that would allow the user to focus energy on phenotypic screening hits that might have novel mechanisms of action and thus avoid compounds with a well-established mechanism or known drug resistance liabilities. In addition, we wished to create a tool that could alert the user to relevant information from work on other species. If a compound is a dihydrofolate reductase inhibitor in humans, it is likely also a dihydrofolate reductase inhibitor in malaria parasites(30).

As described in the methods, this tool, which we have named CACTI consists of 3 modules, coded into python and accepts queries in the forms of tabular or comma-delimited files containing the SMILES of query molecule(s), and additional field columns such as standard name, if desired. The output consists of Excel files from each analysis (module) selected with a comprehensive report for all query molecules mining for synonyms and scholar evidence from the selected knowledge databases including ChEMBL, PubChem, BindingDB and PubMed. Also, a complete list of close analogs to query compounds and their similarities scores from the network construction module, and lastly the clustering module yields a report listing the query compounds and similar molecules from a target set, for example the chemogenomic library, as well as the cluster identification number and percent of similarity between the query molecule and the target molecule. These last two reports can be exported to external tools for network visualizations, such as Cytoscape(31). CACTI can be accessed from the public GitHub repository, and executed by creating a local copy and following instructions indicated in the repository.

### Case study: assessment of drug-target querying using the Pathogen Box dataset

In order to assess the performance of the tool, we applied it to the Pathogen Box dataset(32) from the Medicines for Malaria Venture (**Fig. 2A**). This set is a collection of 400 diverse drug-like molecules, with an average molecular weight of 374.13 Da, active against several neglected tropical diseases. This set was physically assembled by the Medicines for Malaria Venture and shared with users working in the neglected disease space and consists of drug-like starting points for oral drug discovery. Accordingly, only 32 compounds failed to comply the standard Lipinski Rule of Five(33). The set includes 26 positive controls from various scaffold families, 24 of which are part of known drug-target pairs. Controls include well known drugs like the antimalarials doxycycline, primaquine and mefloquine(34), the antibacterial rifampicin(35), the antimicrobials pentamidine, nifurtimox and fluoxetine, the anti-schistosomiasis medicines, praziquantel and mebendazole. For controls, we often found too much information, and these were thus excluded from further consideration. The remaining 374 compounds (**Table S1**) are ones that have reported activity in published phenotypic screens for neglected tropical disease, with the majority being antimalarial or antimycobacterial compounds(36–38) (**Fig. 2B**). Chemical properties for the 400 compounds and biological activity in multiple parasites and pharmacokinetic measurements, including cellular toxicity and pk calculations in HepG2 cells, can be found in ChEMBL-NTD repository (https://chembl.gitbook.io/chembl-ntd) set 21.

**Figure 2.**
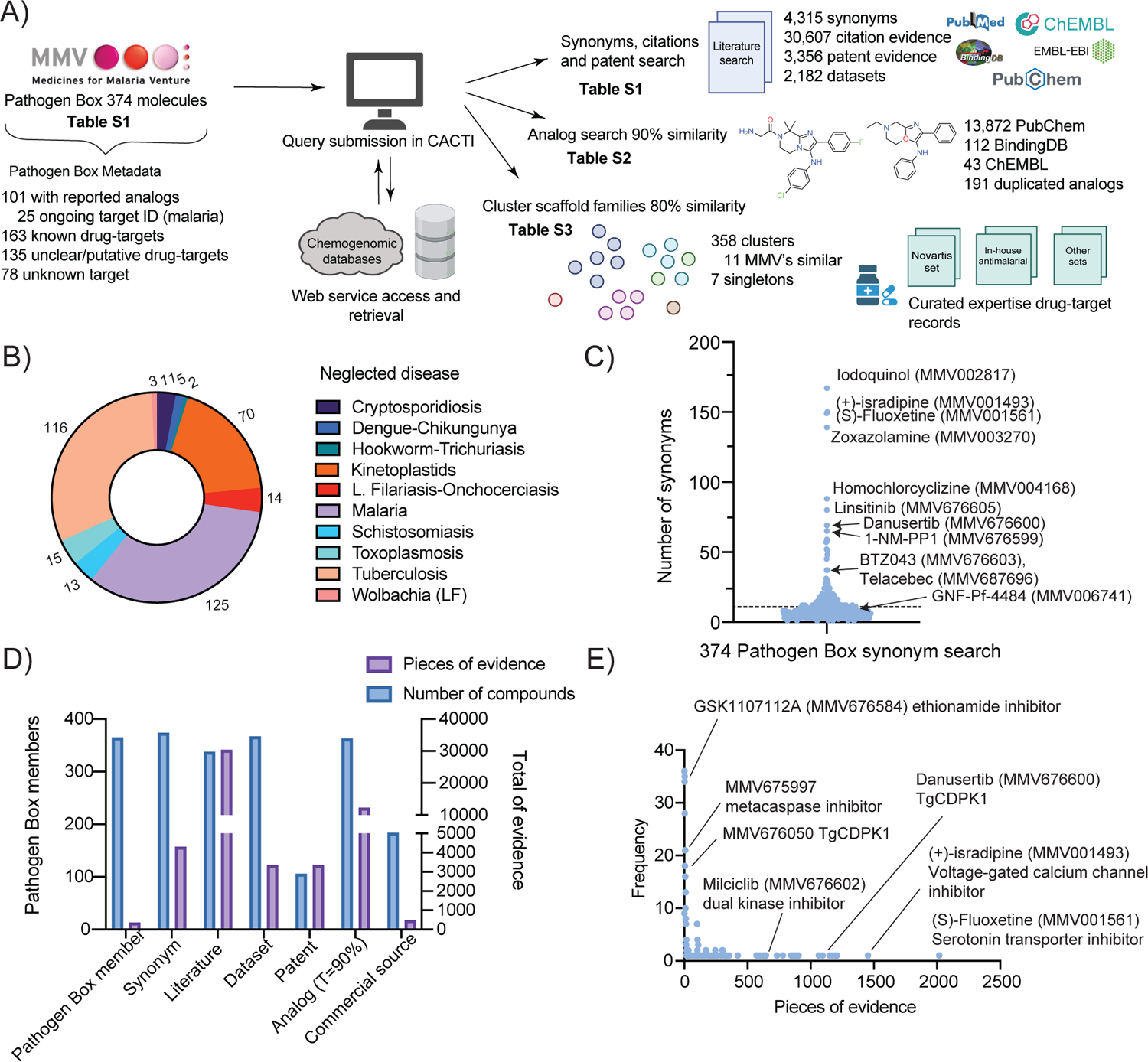
Identification of synonyms, references, and close analogs for 374 Pathogen Box compounds. **A)** Pathogen Box querying workflow. Results obtained of querying 374 Pathogen Box compounds against the pipeline. Supplementary tables related to each analysis are indicated along with the results from each module. **B) Membership of selected compounds.** Number of compounds active against each neglected disease according to the Pathogen Box reference biological data(32). **C) Synonyms identified.** Number of names given for each Pathogen Box compound across different studies. Well established drugs belonging to the Pathogen Box are shown along with the common drug name. A dotted line is shown at the average (11) of synonym names per compound. **D) Statistics for identified information per compound.** Identified categories for the 374 compounds are shown in the X axis. Left Y axis shows the number of Pathogen Box compounds found per category and the right Y axis shows the number of pieces of evidence per category. **E) Pieces of literature found per compound.** Scatter plot showing the total pieces of evidence for each compound (X axis) against the frequency (Y axis). Known inhibitors with their inhibitory protein/enzyme are shown.

### Module 1 identifies 4,315 synonyms for the Pathogen Box compounds

A good literature search can prevent months of wasted effort, but searching biological databases such as PubMed is hampered by the fact that compounds frequently change names in biological publications. We assessed if we could find synonyms for the compounds in an automated way. Searching using CACTI identified 4,315 synonyms that were not included in the initial Pathogen Box description, with an average of 11 names per compound (**Table S1, Fig. 2C-D**). For example, we found 10 synonyms for compound MMV006741, including GNF-Pf-4484, Maybridge3_001644, and BRD-K92165166-001-01-3, among others. Likewise, the automated synonym search revealed that one trivial name for MMV676600 is danusertib, a known pan-Aurora kinase inhibitor(39). Overall, we found 69 different names for MMV676600. These data also provided clues about the provenance of the compounds. The search revealed substantial overlap between the TCAMS library with 164 compounds present and 74 members also present in the St. Jude library, with 74 Pathogen Box compounds existing in both libraries. Additionally, eight Pathogen Box compounds were also found to be members of the Novartis GNF library. These libraries can be accessed at ChEMBL-NTD (https://chembl.gitbook.io/chembl-ntd) set 1 for TCAMS library, set 2 for Novartis, and set 3 for St. Jude set. The automated search also revealed members from the Pathogen Box that are readily accessible to the scientific community. We found 206 compounds with suppliers’ identification numbers for various commercial sources including AKOS, Zinc and MCULE. In general, module 1 was able to capture and identify the naming assigned to Pathogen Box members across studies and public libraries, as well as provide the unique identifiers for selected databases including PubChem (CID identifier) and ChEMBL (CHEMBL identifier).

### Module 1 identifies new citations, datasets and patents for Pathogen Box molecules

No literature citations were provided with the initial Pathogen dataset. Therefore, we also sought to comprehensively identify patent and literature citations using not just the MMV names but all synonyms using CACTI (**Fig. 1E**). Unsurprisingly, searching with the MMV names often produced literature that described the Pathogen Box. However, searching with synonyms provided many more citations on the compounds. For example, MMV675997, is associated with 4 pieces of evidence that were deposited separately under several chemogenomic databases with different synonyms (MMV675997, CHEMBL1094051, BDBM50313769) (**Table S1**). Searching with these synonyms showed that this compound, which has a peptoid backbone and a nitrile P1′ warhead came from a library predicted to contain *T. brucei* Metacaspase enzyme inhibitor(40). Likewise, the kinase inhibitor, danusertib (MMV676600) is associated with a total of 1,195 pieces of evidence including citations indicating a possible role in inhibition of *Toxoplasma gondii* calcium-dependent protein kinase 1 CDPK1 (optimizing small molecule inhibitors of calcium-dependent protein kinase 1), as well as 100 pieces of patented evidence. From these, a CACTI PubMed search using PHA-739358, the additional synonym of MMV676600, yields three more literature references compared to MMV676600 (**Table S1**).

The literature search also showed cases with evidence of similar potent inhibitors across different protozoans. For example, Murphy and colleagues(41) discovered that MMV676050 and MMV676182, both tested against cryptosporidiosis in the Pathogen Box activity profiling, are potent inhibitors targeting CDPK1 in *C. parvum* and *T. gondii* with known crystal structures (PDB ID: 3MWU and 3N51 respectively(41)). Similarly, the literature search revealed that 162 members from the Pathogen Box showed evidence of a drug-target validated in other models, such as cancer cell lines. For example, the chemotherapeutic milciclib (PHA-848125), currently in clinical trials, is shown to be a dual cyclin-dependent and tropomyosin receptor kinase inhibitor (42, 43). Overall, we found 30,425 new pieces of literature, 3,356 patents and 2,182 datasets associated with the Pathogen Box compounds. Though exploring the literature for different names may be straightforward for a small-scale study, as the study size increases, it becomes more difficult to capture these nuances. Therefore, the implemented method of acquiring and cross-referencing all known identifiers to retrieve data from the several chemogenomic databases provides a more detailed and complete report to easily detect further differences.

### Module 2 identifies 12,383 new closely related compounds in the public domain that can be used for SAR by inventory

An important step for compound evaluation and testing is determining if there are closely related analogs available in the public domain that can be sourced for testing. Performing a Tanimoto similarity search for the 374 uncharacterized Pathogen Box members resulted in 12,383 close analogs (*T* = 90%) from the selected chemical knowledge databases (**Table S2, Fig. 2D**). Eleven compounds (MMV393995, MMV676269, MMV676442, MMV021057, MMV202553, MMV688754, MMV099637, MMV676050, MMV676064, MMV676204 and MMV676398) out of the 374 did not report a close analog in the public domain under this criterion. Most analogs identified belonged to the PubChem repository with a total of 12,246, followed by 112 from BindingDB, and 25 identified from ChEMBL database records. On average, analogs resembled the Pathogen Box compounds with a 94.44% similarity, and an average of 34 closely related analogs were identified for each member of the query set. For example, we found 55 analogs of MMV019721, a recently discovered acetyl-coenzyme A synthetase *Plasmodium* inhibitor(44). Many of these analogs are commercially available.

Interestingly, we observed 8 closely related analogs that were identified twice when querying for the 374 Pathogen Box compounds (**Table S2**, **Fig. S2**). Finding a close analog for two different query compounds suggest the presence of similar core scaffolds and likely similar biological targets. For example, the compounds MMV595321 and MMV676477, share five different analogs (CID’s 136636721, 136636828, 156278160, 156278182 and 156285860), most of which are part of an antiparasitic inhibitor optimization effort (patent WO-2021077102-A1). Another analog CID 44526919, related to MMV023969 (TCMDC-134161) and MMV024035 (TCMDC-134227), has shown inhibitory activity against *P. falciparum* up to 97% (IC_50_ < 2 µM)(45) thus the related Pathogen Box compounds could be attractive drug targets.

### Module 3 for new target discovery—guilt by association approaches

For biological function discovery, it is often useful to perform gene expression analysis with the idea that genes with similar expression profiles over many datasets will often have the same function. We reasoned that compounds that are structurally similar might have the same target. To create target predictions, we clustered the 374 Pathogen Box compounds and 156 close analogs having a known target, with sets of 4,716 molecules with known mechanisms of action in order to create hypotheses about their function. The “known target” set was derived from the chemogenetic library assembly by Canham et al.(26) consisting of 4,185 compound-target pairs, 150 known antibacterial drug-target pairs(29), and a collection set of 381 antimalarial compound-target pairs extracted from established databases such as IUPHAR(27) and PDB Ligands(46), in addition to validated ligands originated from phenotypic datasets including TCAMS and GNF Novartis Malaria Box libraries. Though the assembled library of known target pairs was focused on antimalarial inhibitors, many have activity across the parasitic and bacterial diseases included in the study, such as atovaquone and doxycycline, thus we confidently used the set against the Pathogen Box to predict biological functions. Known targets for 152 Pathogen Box compounds (**Table S1**) were incorporated into the prediction analysis. Clustering the 374 and their 156 close analogs with 4,716 known targets resulted in 77 clusters using a similarity threshold of *T* = 80% **(Table S3),** with an average of 4 compounds per cluster, and 264 singletons (**Fig. 3A**). Out of the 77 clusters, twenty-five had two or more Pathogen Box molecules and 71 clusters contained one or more MOA/target or putative target annotations, from which 20 have at least one Pathogen Box compound with unknown or unclear mechanism. We did not find a MOA/target prediction for six clusters (**Fig. 3B**).

**Figure 3.**
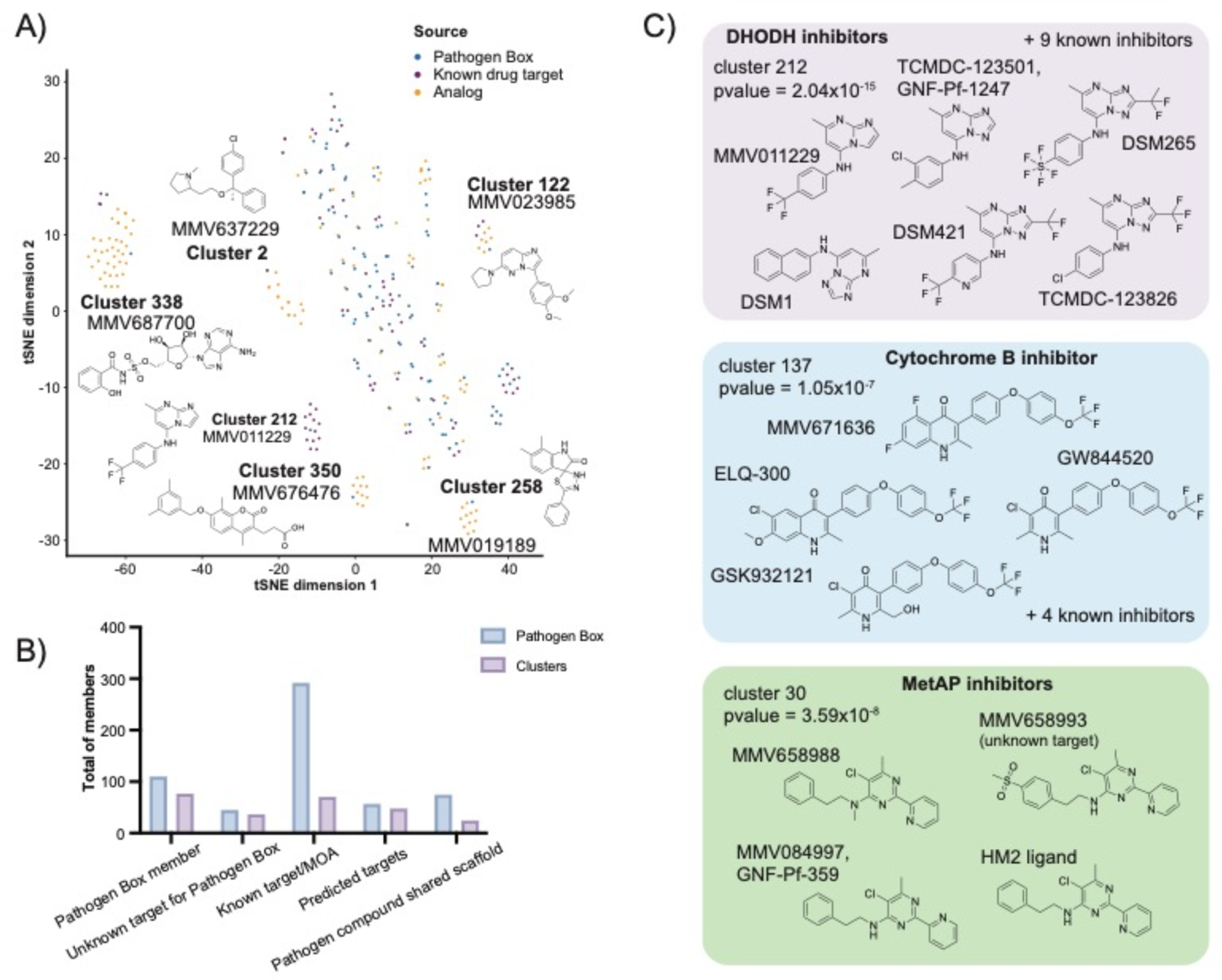
Chemical clustering and target prediction. **A)** Scaffold clustering. Structural similarity calculations with Pathogen Box library and known drug targets, as determined with RDKIT fingerprints. Clusters were assigned using 80% Tanimoto similarity threshold. Visualization of chemical space was performed using scikit-learn t-SNE function. **B) Scaffold clustering statistics.** Total number of Pathogen Box members (light blue) and clusters (light purple) identified for the different categories shown in X axis. Total of known drug-target annotations, and predicted drug-targets, among all clusters are included in the summary categories. **C) Overrepresented clusters with known target.** Examples of clusters with drug-targets found at rates greater than expected according to the hypergeometric mean function. Structures for cluster members are included.

In order to assign targets for the Pathogen Box compounds, we inspected members from each cluster and assigned a potential target/function for the “unknown” Pathogen Box compound based on their counterpart target evidence. From this, we found 71 drug targets from the “known target” set across clusters, of which 75 targets appear repeatedly at rates higher than expected (hypergeometric mean function = 4.4×10^-3^ – 2.6×10^-48^) (**Fig. 3C**). For example, three clusters (55, 56, 212) contain predicted dihydroorotate dehydrogenase (DHODH) inhibitors and six contain predicted cytochrome b inhibitors, of which cluster 62 have one Pathogen Box compound (MMV687807) with no known reported target in *Plasmodium* or *Mycobacterium*. We also found 9 clusters whose members contain an analog or known target set compound from two or more different annotations. This includes cluster 338 where close analogs to MMV687700 are known salicyl-AMP inhibitor and other members from the known drug target set are known human adenosine receptor agonists. Despite this apparent incongruity, all compounds in cluster 338 contain the basic adenosine scaffold and are adenosine analogs. Another example includes cluster 177 where MMV688283 and MMV687246 (PfCDKP5(47)) are clustered together with TCMDC-138293, a known *Plasmodium* DHODH inhibitor. This may suggest some polypharmacology or potential weaknesses in the automated annotations.

We also found cases where the predicted target was assigned based on evidence from a different parasite genus to the one initially tested. The diazine scaffold family (cluster 30) has two compounds that have demonstrated activity against parasite and mammalian methionine aminopeptidase-1b, an enzyme predicted to catalyze the removal of the N-terminal initiator methionine during protein synthesis in parasites and mammals (MMV084997/GNF-Pf-359). The group also contains the uncharacterized anti-kinetoplastid Pathogen Box compounds MMV658993 and MMV658988. This association creates a hypothesis that MMV658993 and MMV658988 may target methionine aminopeptidase-1b in kinetoplasts.

## Discussion

Neglected diseases (parasitic diseases, and to some extent, rare diseases), attract less commercial interest and significantly less overall funding than other diseases, such as cancer or diabetes. On the other hand, much of the small molecule data for well-funded diseases such as cancer lies in the private, well-curated databases that are maintained by pharmaceutical companies. These databases are often one of a company’s biggest intellectual property assets. In contrast, for neglected disease, a larger proportion of the drug discovery data will be generated by academic researchers or public-private partnerships where much of it will ultimately be placed in the public domain. The availability of well curated public screening datasets (e.g. such as the Pathogen Box dataset) creates both opportunities to make new discoveries (e.g. for drug repurposing and target discovery) as well as challenges that relate to the dispersed and disorganized nature of the data. To make the task of assessing diverse data more accessible to neglected disease drug discovery researchers, we constructed a tool to pragmatically assess the major chemical databases and identify all data available for a query compound, as well as similarities between small molecule inhibitors in any system. With this approach a comprehensive report is generated and may be used to redirect laboratory resources in a more focused way, in addition to reducing the amount of work needed for drug-target or chemical optimization efforts.

Our approach has its limitations. First, our algorithm makes no assessment of literature quality. A manuscript reporting that a given compound binds a target may not rigorously assess the quality or strength of the binding. This may happen when a researcher develops a biochemical assay for a target and then tests a library, such as the Pathogen Box library, against this target whereupon the best compounds from this exercise may be reported as potential inhibitors of the target in a publication. Our computational algorithms cannot catch these nuances and are out of scope; although a thorough report on a particular query set will be obtained, a human review component is needed to revise and confirm the veracity/strength of acquired evidence. Perhaps not surprisingly, the data-querying strategy across multiple chemogenomic repositories is especially helpful when querying for small molecules that have been evaluated for activity against multiple disease indications. For example, compounds advancing to clinical-trials or those having a validated drug-target, are query entries that when searched provide the most complete information.

Another limitation is that our approach somewhat relies on the assumption that a compound will have the same target in *T. cruzi* as in *T. gondii*, despite the two species belonging to different eukaryotic phyla. Previous studies have shown that highlight drug-target pairs tend to be conserved across species. A compound like methotrexate likely acts against dihydrofolate reductase in human and in malaria parasites. Cladosporin targets lysyl-tRNA synthetase in both yeast and malaria parasites. There are hundreds of other examples of target conservation across species. Species selectivity thus comes from natural or engineered specificity as well as innate ability of different species to detoxify compounds. On the other hand, just because a compound targets the electron transport chain in malaria parasites doesn’t mean that it targets the electron transport chain in *M. tuberculosis* (e.g. such as in ELQ-300, with unreported target in tuberculosis) and predictions and caution is needed.

Another concern is that we relied on chemical scaffold clustering: A chemical comparison depends on the chemical fragment partition that is selected, resulting in potential arbitrary cluster assignment. In addition, clustering is less suitable for natural products, which tend to have more complex structures than smaller molecules. Increased size means partition methods favor fragment partitions that break down molecules and ignore the R groups that may be vital for the pharmacological effect of the natural product. Thus, although the provided scholarly report will be helpful, the clustering approach as scripted is less suitable for their evaluation. Nevertheless, we found that close analogs with similar drug-target matches did tend to group together even if the larger group often split into more than one cluster. We are confident that the established prediction method provides an initial hint for further target validation efforts.

A limitation with this approach includes the predictive component of the analysis. Though is accepted that similar molecules tend to have similar mechanisms of action or drug-targets, experimental validation is needed to confirm similar inhibitory potency and binding to the desired target. The assumption that one small molecule will have just one target is more fraught as compounds get larger and where two different chemotypes (e.g. an ATP-like molecule and a naphthalene) may be incorporated into one small molecule. In addition, for promiscuous inhibitors, like kinase inhibitors, our approach may provide limited resolution.

Another concern is that one component of this tool relies completely on network connectivity and database server availability (e.g. open query-request for selected databases) in order to query and retrieve literature and analogs from the public domain. To partially address this need, a future avenue includes the use of a periodically “database dump” that can be locally accessed and queried in lieu of internet access when connectivity is unreliable or inaccessible.

Overall, we believe the identification of literature evidence and close analog search obtained through CACTI will be a valuable resource that will aid researchers in early stages of drug discovery pipeline, to consequently reduce the exploration space in a more focused set of small molecules for further quantitative structure-activity relationship analysis. Similarly, the target prediction module implemented in this tool will serve as starting points for subsequent drug-target validation efforts and support the translation of genomic data into effective new drugs through the comparison of scaffold families. This automated tool shows a promising approach to quickly investigate multiple chemical entities in a single query, and prioritize hits for further exploration, especially in academic settings where compound management and resupply is limited.

## Data Availability

CACTI is accessible via GitHub (https://github.com/winzeler-lab/CACTI) for download and personal use.

## Acknowledgements

This work was funded by a grant from the Bill and Melinda Gates Foundation (INV-039628). The authors thank MMV for their contribution of the Pathogen Box collection and making it available to the community.

## Author Contributions

E.A.W. conceived experiments, provided funding, provided supervision, wrote the manuscript, reviewed the manuscript, constructed supplemental files, performed analysis and designed figures. K.P.G.-M. wrote the first draft of the manuscript, reviewed the manuscript, constructed supplemental files, performed analysis and constructed figures.

