## Supplementary Material for "*CACTI: An in-silico* drug-target prediction tool through the integration of chemogenomic data and clustering analysis"

Affiliations
[1] Department of Pediatrics, University of California, San Diego, School of Medicine, La Jolla, CA 92093 USA

**Supplementary Table S1. Pathogen Box description, synonyms and scholar evidence found**. Attached Excel spreadsheet.

**Supplementary Table S2. Close analogs to Pathogen Box compounds.** Attached Excel spreadsheet.

**Supplementary Table S3. Cluster analysis against external sets.** Attached Excel spreadsheet.

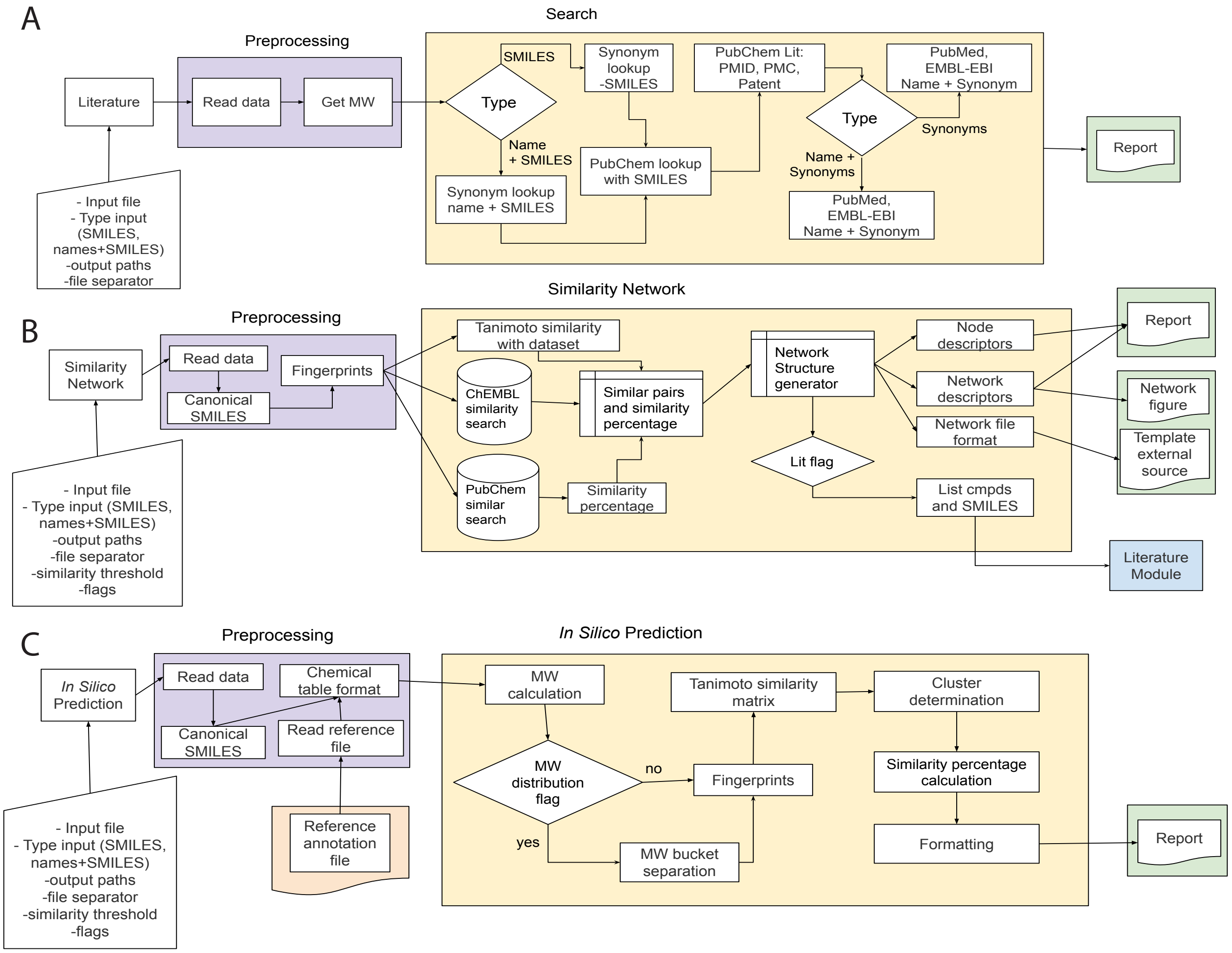

**Supplementary Figure S1. Modules for drug target prediction.** Module workflow for **A)** literature search, **B)** similarity network and analog search, and **C)** *in silico* prediction. Trapezoidal boxes show input data types of choice for each module, purple boxes represent preprocessing steps to each section. Yellow boxes delimitate the steps to complete a module's task, where diamonds are decision steps and rectangles a function to be performed. Green boxes are figure or results report generated.

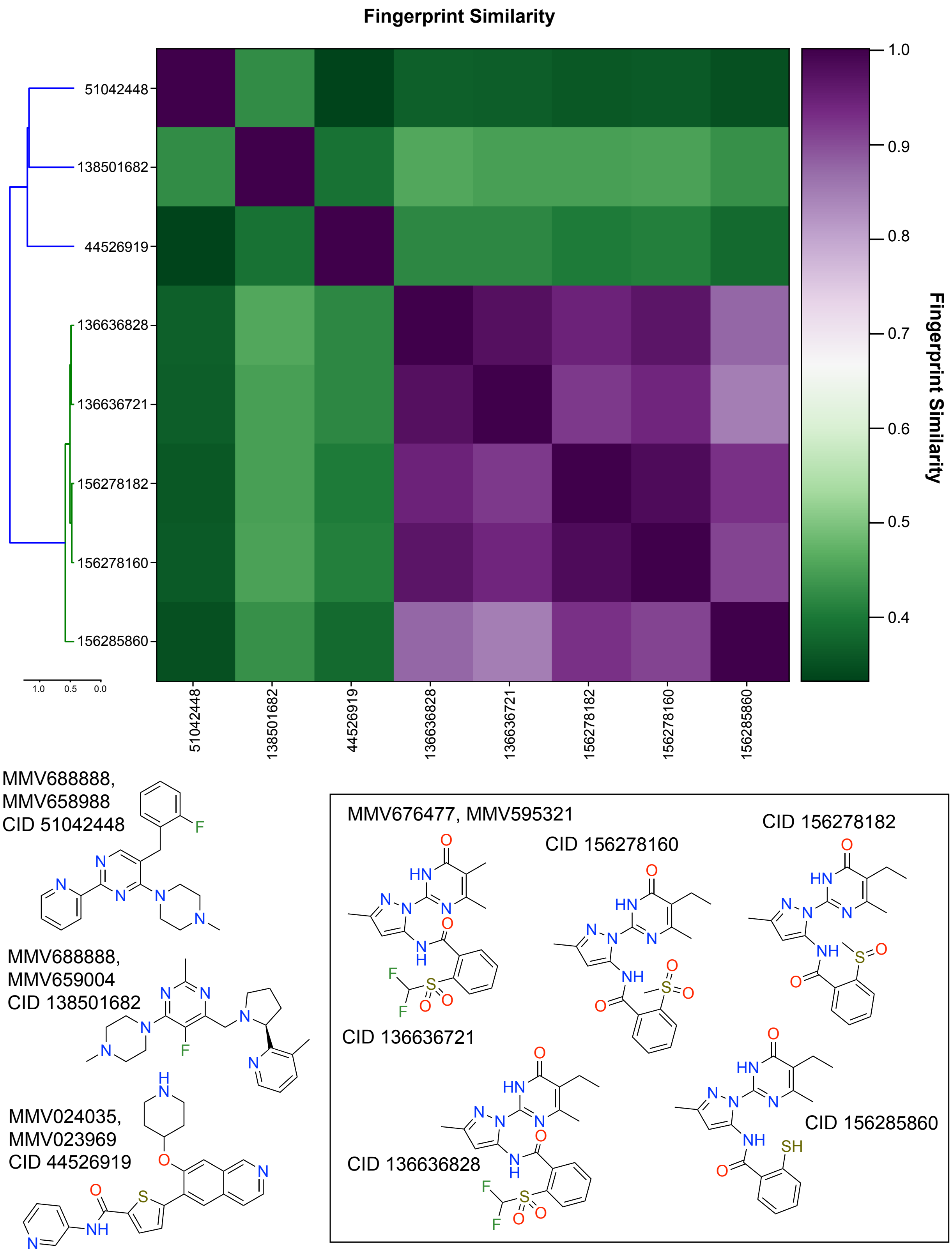

**Supplementary Figure S2. PubChem duplicated analogs fingerprint similarity.** Scaffold similarity comparison for the 8 PubChem (CID) close analogs for two different Pathogen Box query compounds. Fingerprint similarity scores were calculated using RDKIT[1] Tanimoto similarity function. Left panel shows the hierarchical clustering for each CID based on the calculated similarity, central panel shows the similarity intensity color coded according to the left panel gradient, where low similarity (0-0.5) is represented by green and high similarity (0.5-1) is represented by purple. Structures for analogs are shown, including the source Pathogen compound.

**References**

1 RDKit. *RDKit: Open-source cheminformatics*, <https://www.rdkit.org>
